# BlackSheep: A Bioconductor and Bioconda package for differential extreme value analysis

**DOI:** 10.1101/825067

**Authors:** Lili Blumenberg, Emily Kawaler, MacIntosh Cornwell, Shaleigh Smith, Kelly Ruggles, David Fenyö

## Abstract

Unbiased assays such as shotgun proteomics and RNA-seq provide high-resolution molecular characterization of tumors. These assays measure molecules with highly varied distributions, making interpretation and hypothesis testing challenging. Samples with the most extreme measurements for a molecule can reveal the most interesting biological insights, yet are often excluded from analysis. Furthermore, rare disease subtypes are, by definition, underrepresented in cancer cohorts. To provide a strategy for identifying molecules aberrantly enriched in small sample cohorts, we present BlackSheep--a package for non-parametric description and differential analysis of genome-wide data, available at https://github.com/ruggleslab/blackSheep. BlackSheep is a complementary tool to other differential expression analysis methods that may be underpowered when analyzing small subgroups in a larger cohort.

## Introduction

Proteogenomic studies characterizing cancer have been completed by many groups, several of which also conducted phosphoproteome analysis (1–7). Outlier identification was used in a number of these studies to identify samples with aberrantly high levels of each phosphosite and phosphoprotein (1,8). In these studies, outlier identification and subsequent subtype enrichment was used to interpret phosphopeptide data at the protein-level, and highlight novel putative clinically relevant targets (1) or to nominate targets in a kinase inhibitor screen for sensitizers in drug-resistant cell lines (8). This non-parametric method is of particular use for multi-omics studies, as non-parametric approaches are more robust to the various sources of technical noise present in these data sets, which violate assumptions in parametric tests.

Outlier values in a dataset are often assumed to be experimental artifacts and are discarded prior to downstream statistical analyses. However, sometimes recurrent outliers are the most meaningful values in a dataset, representing profound biological effects. In particular, when characterizing biological systems and identifying disease vulnerabilities, the largest changes in abundance are often the most revealing (9,10). Furthermore, many diseases, including cancer, are heterogeneous, with significant molecular variability requiring highly personalized approaches for successful treatment. Current strategies for identifying characteristic molecular patterns for groups of samples are underpowered for rare disease subtypes and use assumptions about the underlying distributions of the features in question, which are often inaccurate and/or discard extreme values with biological significance. We propose a complementary strategy using the enrichment of outlier values within subtypes for characterizing disease subtypes, informing diagnostic panels, and potentially designing personalized therapeutic strategies for individual patients.

## Materials and Methods

BlackSheep is an easy-to-use package available on Bioconductor (https://github.com/ruggleslab/blacksheepr) and Bioconda (https://github.com/ruggleslab/blackSheep). It can be used in R, python or as a command line utility. BlackSheep has two major components: the ‘DEVA’ (<u>D</u>ifferential <u>E</u>xtreme <u>V</u>alue <u>A</u>nalysis) module for calling outliers, collapsing features by parent molecule (i.e. phosphopeptides to a protein) and differential analysis, and the ‘run_simulations’ module for assigning p-values to each outlier call. The input data is an expression matrix, structured as rows of features (genes, proteins, phosphosites, etc.) and sample columns, and a sample annotation file used to group samples for comparisons (**Supplementary Table 1A, 1B**). No prefiltering is necessary or recommended for DEVA. Normalization of the input matrix is strongly recommended; a function for this is provided. For normalized data, we suggest a sample coverage normalization followed by log_2_ transformation.

### Differential Extreme Value Analysis (DEVA)

To call outliers, the median and interquartile range (IQR) for each row is calculated. The user specifies whether to call overly abundant (i.e. up) or depleted (i.e. down) values. Outliers are defined as any value more than a multiple of the IQR above or below the median, where the multiple of the IQR is user-specified, with a default of 1.5 (**Fig. 1A)**. After calling outliers, there is an optional aggregation step for collapsing rows containing related features into a single row (e.g. many phosphosites collapsed into a protein). Aggregation is achieved by counting outliers and non-outliers separately for each protein. The output is two tables, one with outlier and non-outlier counts per protein used for downstream comparisons (**Supplementary Table 2A, 2B**), and the other containing the fraction of outliers in each sample which is helpful for visualization (**Supplementary Table 2C, 2D**).

**Figure 1.**
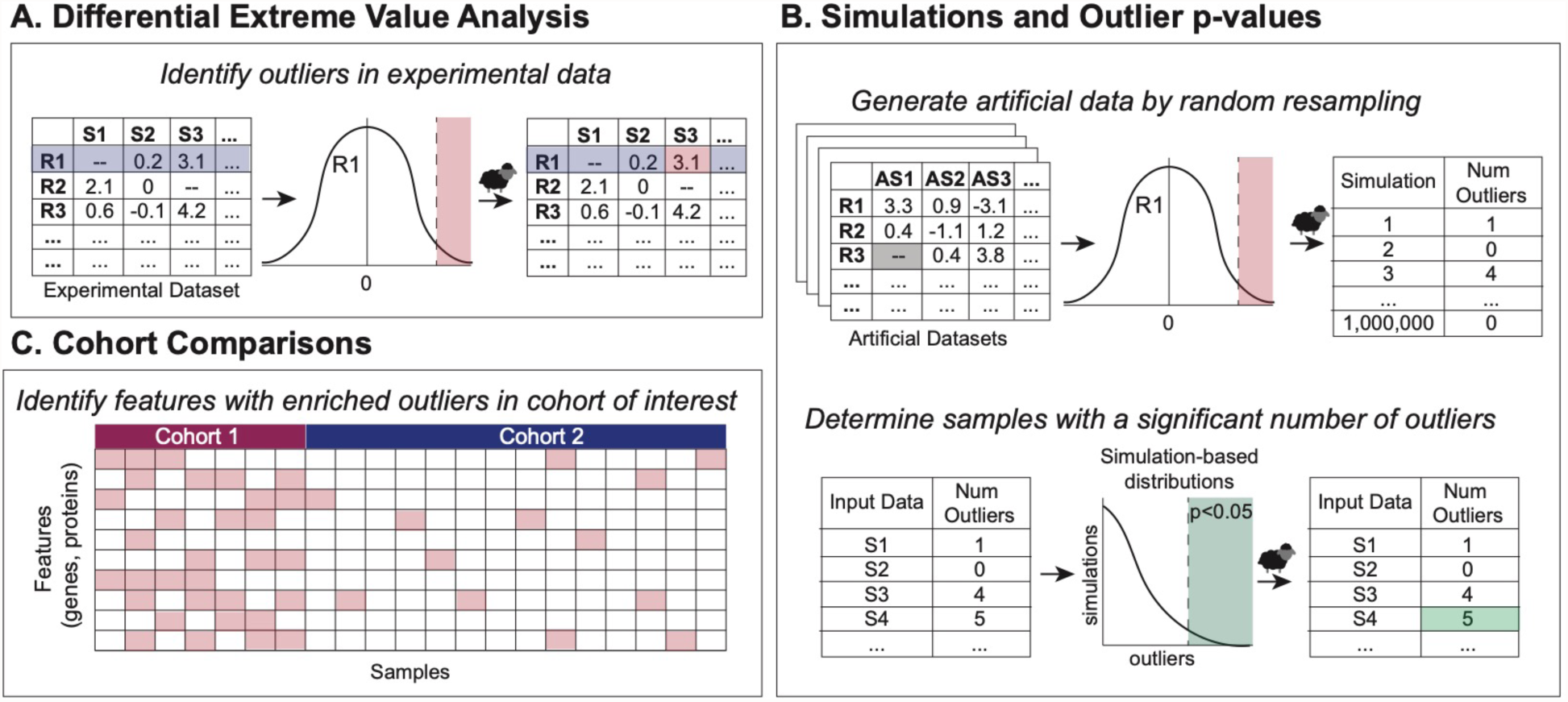
BlackSheep Workflow. (A) Outliers are identified for each feature (row) in the experimental dataset and (B) using simulations and data resampling significance value assigned for each sample/feature. (C) Cohort comparisons identify features with enriched outliers within a sample cohort of interest.

### Simulations and outlier p-values

The second main function in the package is ‘run_simulations’, which uses simulations based on the observed data to calculate a p-value for each sample for each parent molecule. For each repetition (“simulated sample”), the procedure is as follows. First, for each feature in the parent molecule (e.g. phosphosite on a protein), its value in a simulated sample is determined to be either observed or missing; the likelihood is based on the proportion of missing values for that feature in the actual data. Next, if its value is determined to be “observed”, a random value is assigned from a Gaussian KDE fit to the observed values from the associated feature. The assigned value is tested against the outlier threshold for that feature to determine outlier status. This is repeated for all features related to the parent molecule. The frequency of outliers found in the simulated data is used to assign a p-value to the observed data based on the number of outliers found for each parent molecule in each observed sample. A significance threshold is set at a user defined alpha (e.g. p<0.10). The output file (**Supplementary Table 3**) contains a p-value for outlier status for each parent molecule in each sample if it reaches significance (**Fig. 1B**).

### Cohort comparisons

Groups of samples can be compared with DEVA to identify features with enrichment of outliers within a group. For every comparison in a user-supplied annotation table (**Supplementary Table 1B**), BlackSheep calculates enrichment of outliers for every group of samples identified in the annotation table. Analysis can be limited to a user-supplied list of genes, such as kinases (1).

To calculate enrichment, first a row-based filter is applied, removing rows where the average rate of outliers is lower in the group of interest than in the outgroup. Second, to ensure that results are not driven by a small subset of the group of interest, we only keep rows that have at least one outlier value in a user-defined proportion of samples in the group of interest; the proportion defaults to 0.3. Finally, DEVA performs a Fisher’s exact test on counts from outlier and non-outlier values in the group of interest vs the outgroup. All p-values are then corrected for multiple hypothesis testing using the Benjamini-Hochberg procedure. Results can be output as a table of q-values for all comparisons (**Supplementary Table 2E, 2F**); a table with outlier counts, p-values, and q-values per comparison (**Supplementary Table 2G, 2H**); or a heatmap showing values in each sample for rows with significant enrichments of outliers (**Fig. 2B**).

**Figure 2.**
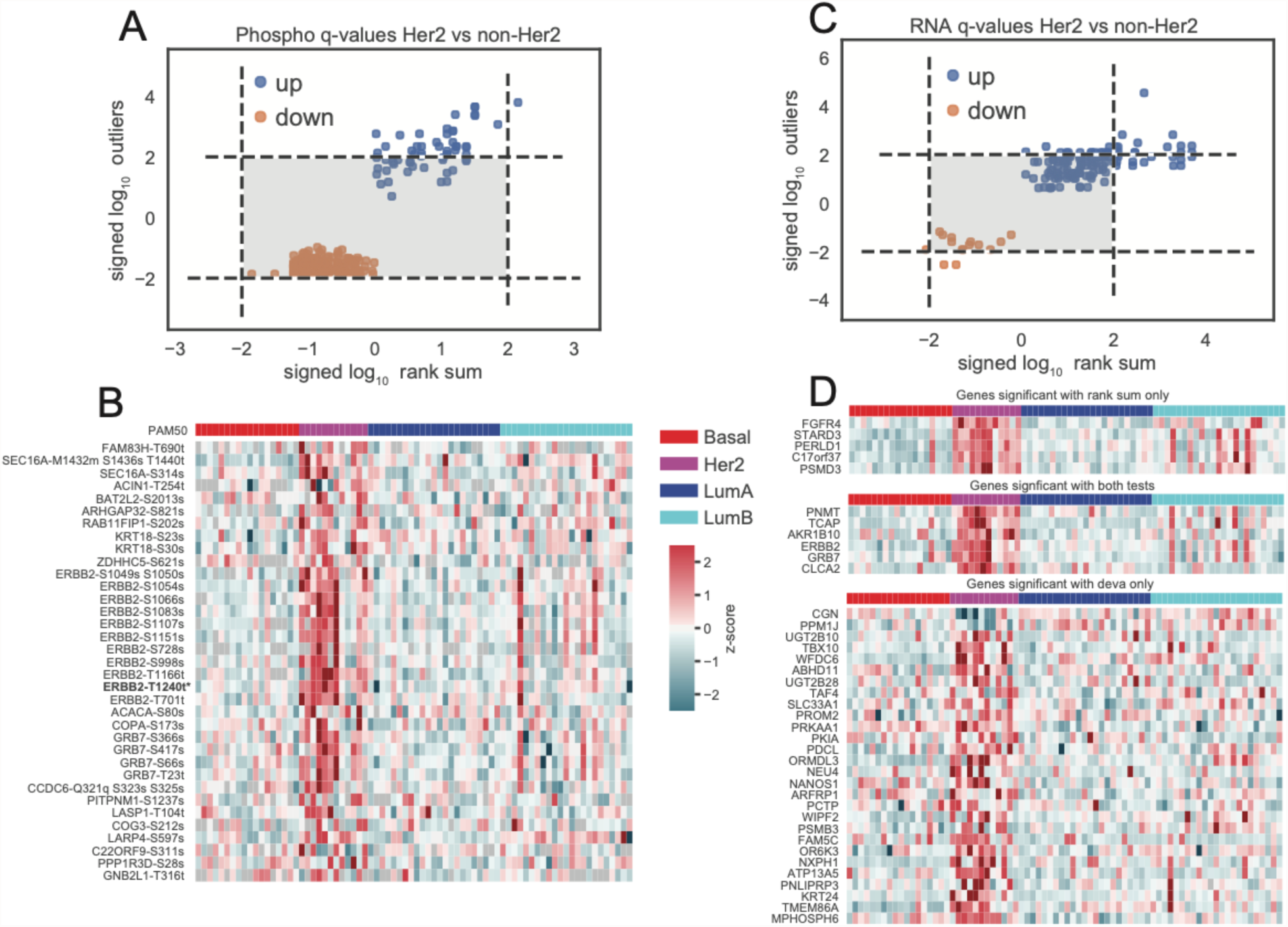
Comparing BlackSheep and Rank-Sum Tests. (A, C) Signed log_10_ q-values from blacksheep.deva and rank-sum tests when comparing normalized values in Her2e vs all other samples in (A) phospho and (C) RNA data. Dotted lines indication FDR < 0.01. (B) Z-scores of relative log_2_ abundance of all phosphosites with FDR < 0.01 calculated by BlackSheep. * indicates ERBB2-T1240 had FDR < 0.01 using a rank-sum test. (D) Z-scores of log2 relative abundance of RNA with FDR < 0.01 calculated by rank-sum only (top), both methods (middle) or blacksheep.deva only (bottom).

## Results

### Application to breast cancer cohort

Reproducibly hyperphosphorylated kinases within a specific subtype or patient cohort represent attractive targets for future drug development and repurposing (13–18). To demonstrate the utility of BlackSheep, we applied it to a dataset from a proteogenomic breast cancer study (1) to find putatively over-active kinases, unique to a molecular subtype (1,11,12). Here, we compare the results of BlackSheep to the commonly used rank-sum test to identify differentially abundant phosphosites in Her2-enriched (Her2e) breast cancer samples vs all other samples (**Fig. 2A**). In this cohort, Her2e is the smallest group, comprising 12 of the 76 samples. Using the full phosphosite expression matrix (63,130 phosphosites, 9881 proteins), results of BlackSheep and the rank-sum test were corrected for multiple hypotheses with equal stringency. At an FDR cutoff of 0.01, rank-sum identified one enriched phosphosite, on the Her2 (ERBB2) protein: ERBB2-T1240 (**Fig. 2A**); the DEVA pipeline identified 10 additional phosphosites on ERBB2, as well as phosphosites on established co-amplicons and modulators of Her2 signaling, such as GRB7 (19,20) (**Fig. 2A-B**). When applied to RNA abundance data from the same cohort, BlackSheep and rank-sum tests identify many of the same enriched genes at FDR < 0.01 (**Fig. 2C, 2D**). In addition, rank-sum found genes that have high values in the group of interest and samples in the outgroup (**Fig. 2D, top**) while BlackSheep identified additional genes that are exclusively enriched or depleted in only the group of interest (**Fig. 2D, bottom**). BlackSheep does not identify features that are enriched in large fractions of samples within a cohort – a feature with consistently high or low values in a large fraction of the cohort will increase the median and IQR, and will no longer be called an outlier. For small groups within a cohort, BlackSheep is able to identify enriched features (**Fig. 2A**).

## Conclusion

Several cancer types have patients that fall into rare subgroups with worse prognoses than the majority of patients (e.g. serous in endometrial cancer, basal-like in breast cancer). Due to the difficulty in acquiring sufficient numbers of samples, these patients are the hardest to study, yet they are the patients most in need of new therapies. While standard analysis techniques are useful for finding characteristics that are enriched in large subgroups of samples, these strategies often lack the power to find the same for small subgroups. BlackSheep provides a user-friendly, complementary method to delineate enrichments in a small group of samples within a cohort. We show that BlackSheep can find enrichment of known markers for small groups of samples, such as ERBB2 and GRB7 in Her2e samples, which other commonly used analysis paradigms miss. BlackSheep is a flexible complement to other methods such as rank-sum tests. In the future, BlackSheep-like strategies can be applied in the clinic to design and interpret diagnostic panels applied to single tumors, to highlight targets of drugs that can be repurposed for new indications, and the “run_simulations” module can be particularly useful to devise personalized treatments by prioritizing drugs that target significant outliers in a tumor.

## Tables

**Supplementary Table 1. Example Input Files.** (A) Data expression matrix, structured as rows of features (genes, proteins, phosphosites, etc.) and sample columns, and (B) sample annotation file, containing comparison group labels for each sample.

**Supplementary Table 2. Example Output Files from DEVA.** (A, B) Outliers output matrix containing the number of (A) up and (B) down outliers per row per sample, (C, D) the fracTable matrix containing the fraction of rows mapping to that parent molecule (e.g. gene) with (C) up and (D) down outliers per sample. (E, F) a significance output file containing a q-value for each row passing this adjustable percentile filter in any comparison for (E) up and (F) down outliers. (G, H) a table of outlier counts, p-values and q-values for the Her2 comparison in (G) up and (H) down outliers.

**Supplementary Table 3. Example Output from run_simulations.** (A) p-values associated with each parent molecule (e.g. gene) for each sample.

## Acknowledgements

This work has used computing resources at the NYU High Performance Computing Facility (HPCF).

